# Comparative transcriptomic analysis of intestinal Tuft cells reveals species selectivity of DCLK1 expression

**DOI:** 10.64898/2026.10.29.752682

**Authors:** Abbey J. Kirkwood, Karly E. Masse, Van B. Lu

**Affiliations:** Department of Physiology and Pharmacology, University of Western Ontario, London, ON, Canada; Children’s Health Research Institute, London Health Sciences Centre, London, Ontario, Canada

**Author notes:** Corresponding author: V.B. Lu.

## Abstract

Intestinal tuft cells are an emerging class of cells under investigation for its potential role in colorectal cancer and inflammatory bowel disease. These cells are a rare population of cells within the intestinal epithelium, thus requiring the use of cell type markers to identify tuft cells for study. Mouse models have primarily targeted the *Dclk1* gene to identify tuft cells; however, in human studies it has been more challenging to identify *DCLK1* expression in the intestine. This rapid report aims to provide clarity on appropriate cell type markers to identify human intestinal tuft cells for study.

## Manuscript

Intestinal tuft cells were first described based on their distinctive morphology, characterized by a prominent apical tuft of microvilli that extends into the intestinal lumen. Subsequent studies identified doublecortin-like kinase 1 (DCLK1 or DCAMKL1) as being highly expressed in intestinal tuft cells, establishing DCLK1 as a widely used marker for their identification in mice^1,2^. The ability to readily identify and genetically target DCLK1-expressing cells has facilitated extensive investigation of tuft cell biology in murine models, including their role in the epithelial “weep and sweep” response to helminth infection and the initiation of associated type 2 immune responses^3–5^.

In humans, however, the identification and study of intestinal tuft cells has been considerably more challenging. Although DCLK1 has been identified in human intestinal tissue, studies have shown that DCLK1-expressing cells are morphologically more like enterocytes, and that human tuft cells might not express DCLK1 at all^6^. Several alternative markers, including choline acetyltransferase (CHAT), advillin (AVIL), and KIT proto-oncogene, receptor tyrosine kinase (KIT) have subsequently been associated with human tuft cells^7^. However, none have emerged as a sufficiently specific and broadly applicable marker for identifying human intestinal tuft cells in isolation. Consequently, studies of human tuft cells often require prior identification or enrichment of the epithelial population, limiting their utility for direct identification and targeting of tuft cells.

To address this limitation, we re-analyzed publicly available single-cell and bulk-tissue RNA sequencing datasets from mouse and human intestine to define conserved tuft cell transcriptional signatures and identify candidate markers with preferential expression in human tuft cells.

We first characterized mouse intestinal tuft cells using publicly available intestinal epithelial scRNA-seq datasets (GSE92332, GSE245315, and GSE251694) and bulk-tissue RNA-seq datasets (GSE262198 and GSE114067). Mouse tuft cells exhibited the expected expression of established tuft cell-associated genes, including *Dclk1, Sh2d6, Avil, and Trpm5*, consistent with previously reported transcriptional profiles (Figure 1A+B).

**Figure 1.**
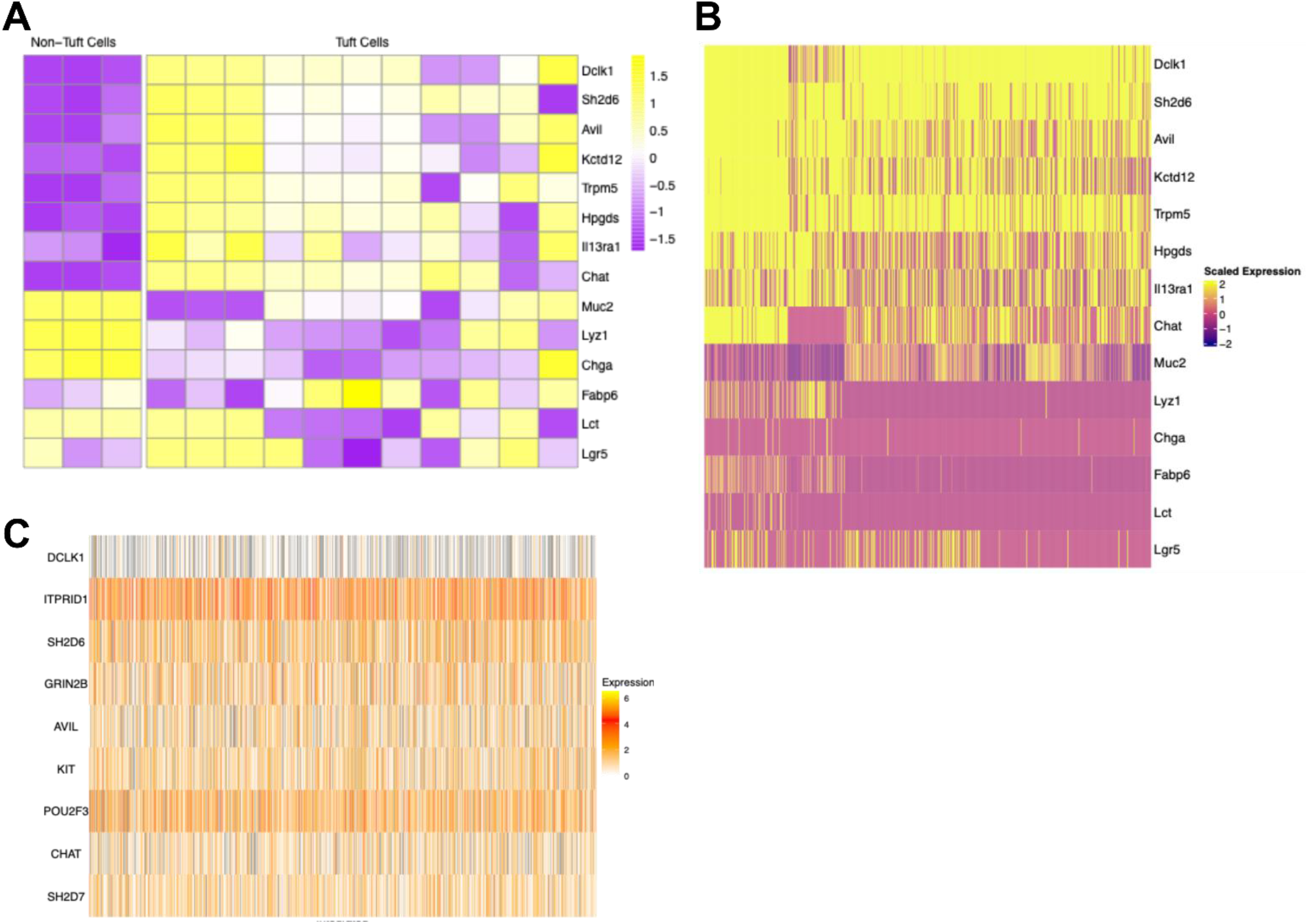
Heatmap representation of transcriptomic analysis of mouse and human intestinal tuft cells. **(A)** Bulk-tissue RNA sequencing heatmap showing mouse tuft cell marker expression across 2 independent datasets. Each box represents a distinct biological replicate, with yellow indicating high expression, and purple indicating low expression. **(B)** Single-cell RNA sequencing heatmap showing mouse tuft cell marker expression across 3 independent datasets. Each line represents a single cell, with yellow representing high expression, and purple indicating low expression. **(C)** Single-cell RNA sequencing heatmap showing human tuft cell marker expression. Each line represents a single cell, with yellow representing high expression, and white or grey representing low or no expression, respectively.

We next compared the expression of established tuft cell-associated genes across mouse and human datasets to identify transcriptional features that were conserved between species. *POU2F3, AVIL, KIT*, and *CHAT* were consistently expressed in the human tuft cell population, with *POU2F3* demonstrating the most uniform expression. In contrast, *DCLK1* showed strong enrichment in mouse tuft cells but minimal expression in human tuft cells (Figure 1C).

Immunofluorescence experiments confirmed that DCLK1 is not expressed in human intestinal tissue, whereas it was successful in identifying tuft cells in the mouse tissue (Figure 2). We were able to confirm the presence of tuft cells in human intestinal tissue by positive IL-25 immunoreactivity, which is the primary transmitter exclusively released from tuft cells (Figure 2). These findings were consistent with our transcriptomic analysis and further demonstrated that the species-specific differences in DCLK1 expression observed at the transcriptional level are reflected at the protein level.

**Figure 2.**
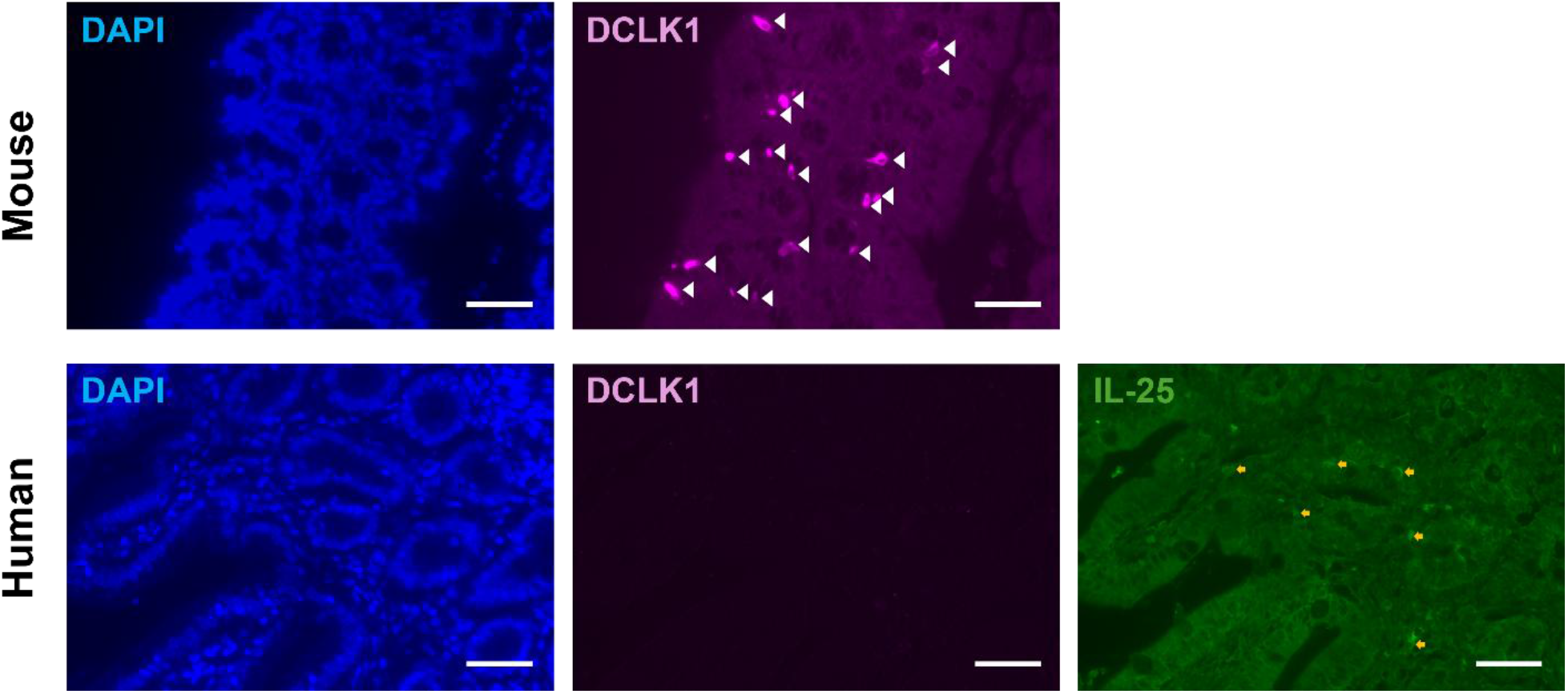
Differential DCLK1 expression in mouse and human intestinal tissue. Representative immunofluorescence images of mouse (top) and human (bottom) intestinal tissue. Nuclei were stained with DAPI (blue), DCLK1 immunoreactivity shown in magenta with cells indicated by white triangles, and IL-25 immunoreactivity shown in green with cells indicated by orange arrows. Scale bars = 10 μm.

Together, these analyses reveal a marked species difference in the transcriptional identity of intestinal tuft cells. While *Dclk1* represents a prominent feature of the murine tuft cell transcriptional program, its expression is substantially reduced or absent in human tuft cells. Conversely, *POU2F3* and other tuft cell-associated genes are retained in the human tuft cell population, suggesting that tuft cell identity is conserved despite substantial differences in the expression of individual canonical markers.

These findings have important implications for the study of human intestinal tuft cells. Much of our current understanding of tuft cell biology has been derived from murine models in which DCLK1 is used as a marker and genetic tool for tuft cell identification. Our data demonstrate that this feature is not conserved in human intestine, highlighting an important limitation when translating DCLK1-based approaches from mouse to human systems. The identification of conserved transcriptional features, particularly POU2F3, may therefore provide a more appropriate framework for defining human tuft cell populations.

More broadly, these findings underscore the importance of validating cell-type markers across species rather than assuming that markers established in model organisms retain equivalent specificity in human tissues. Comparative transcriptomic approaches may be particularly valuable for identifying conserved cellular programs while distinguishing species-specific features that could influence the interpretation of human intestinal biology.

Together, our transcriptomic and experimental analyses demonstrate a clear species-specific difference in DCLK1 expression in intestinal tuft cells. While *Dclk1* is strongly associated with murine tuft cells and effectively identifies this population in the mouse intestine, *DCLK1* expression is low or absent across most human tuft cells and therefore does not provide an equivalent marker in human intestine. In contrast, *POU2F3* and other tuft cell-associated genes were consistently expressed in the human tuft cell population, suggesting that core features of tuft cell identity are conserved despite differences in individual marker expression. These findings highlight an important consideration for translating tuft cell biology from mouse models to human systems and emphasize the need to validate cellular markers across species. Establishing markers that accurately identify human tuft cells will be critical for their isolation, characterization, and functional study in human intestinal models.

## Methods

### Transcriptomic analysis of mouse bulk RNA-seq datasets

Raw bulk RNA-sequencing datasets (GSE262198^8^, GSE114067^9^) were obtained from the Sequence Read Archive (SRA) and downloaded using the SRA Toolkit. SRA files were converted to FASTQ format and reads were quality controlled and trimmed by fastp. Reads were trimmed from the 5’ and 3’ ends based on a mean quality threshold of 20 and reads shorter than 30 bp following trimming were removed. For sequence alignment, the *Mus musculus* GRCm39 primary assembly reference genome was obtained from Ensembl release 109, and a genome index was generated using HISAT2 (v2.2.1). Reads were aligned to the reference genome, and the resulting alignments were converted to BAM format and sorted using SAMtools (v1.18). Gene-level read counts were generated from the sorted BAM files using featureCounts.

For downstream analysis, the gene-level count matrix was imported into R and processed using DESeq2 (v1.48.2). Genes with a mean normalized read count greater than 10 across samples were retained and variance-stabilizing transformation was performed for visualization. A panel of marker genes representing tuft cells and other intestinal epithelial cell populations was selected and expression values for these genes were extracted from the matrices. The selected genes were standardized across samples by calculating row-wise z-scores, allowing relative expression patterns between samples to be visualized. Heatmaps were generated with the pheatmap (v1.0.13) package.

### Transcriptomic analysis of mouse intestinal scRNA-seq datasets

Single-cell RNA sequencing data from three published datasets (GSE245315^10^, GSE251694^11^, GSE92332^12^) were processed using the Seurat (v5.4.0) package in R (v4.5.1). Cells with at least 200 detected genes were retained, and genes detected in at least 3 cells were included in the initial Seurat objects. After filtering out low-quality cells based on the number of detected genes, total RNA counts, and mitochondrial gene expression, cells with fewer than 1000 or more than 5000 detected genes, more than 20 000 total RNA counts, or greater than 25% mitochondrial gene expression were excluded. The remaining data were normalized, and highly variable features were identified using the Seurat package.

The filtered datasets were subsequently integrated across biological replicates using Seurat’s anchor-based integration workflow. The integrated assay was used for downstream analysis, and the data were scaled prior to dimensionality reduction. Principal component analysis (PCA) was performed using the first 30 principal components, and cells were clustered using a resolution parameter of 0.5. The tuft cell population was identified based on the expression of *Dclk1*, visualized using the FeaturePlot function. Marker genes associated with the identified tuft cell cluster were subsequently determined using the FindMarkers function. Heatmaps were generated with the ggplot2 (v4.0.1) package.

### Transcriptomic analysis of human intestinal scRNA-seq dataset

Single-cell RNA sequencing data were processed separately for each region of the intestine from published dataset HBM692.JRZB.356^13^ using the Seurat (v5.4.0) package in R (v4.5.1). Raw gene expression matrices were imported and Seurat objects were generated for each sample by retaining genes detected in at least 3 cells and cells with at least 200 detected features. Cells were filtered based on quality-control metrics, retaining cells with greater than 200 and fewer than 6,000 detected genes and less than 15% mitochondrial gene expression. The remaining data were normalized, and highly variable features were identified using the Seurat package. Samples were integrated using Seurat’s anchor-based integration workflow with reciprocal PCA (RPCA) as the dimensionality reduction method. The integrated assay was used for downstream analysis, and the data were scaled prior to dimensionality reduction. Principal component analysis (PCA) was performed using the first 30 principal components and cells were clustered using a resolution parameter of 0.5. The tuft cell population was identified based on the expression of previously reported human tuft cell markers or lineage-defining genes (*POU2F3, TRPM5, IL25, AVIL*, and *KIT*) and visualized using the FeaturePlot function. Marker genes associated with the identified tuft cell cluster were subsequently determined using the FindMarkers function. Tuft cell clusters identified in each object were used to examine the expression of tuft cell-associated genes across each intestinal region. Heatmaps were generated with the ggplot2 (v4.0.1) package.

### Animal tissues and tissue processing

Animal studies were approved by the Western University Animal Care Committee according to guidelines established by the Canadian Council on Animal Care. Intestinal tissue was collected from female C57BL/6 mice aged 2-3 months following euthanization. Intestinal contents were flushed with ice-cold phosphate-buffered saline (PBS) and rolled longitudinally into a “Swiss-roll” as previously described^14^. Briefly, tissues were fixed overnight at 4ºC in 4% paraformaldehyde (PFA; Electron Microscopy Science 15713-S) in PBS. The fixed tissues were then rinsed with PBS, cleared with xylene substitute, and rehydrated using a series of graded ethanol solutions. Paraffin wax was used to embed tissues overnight. The formalin-fixed paraffin-embedded blocks were serially sectioned used a rotary microtome (Leica Biosystems HistoCore MULTICUT) at a thickness of 5 μm. Sections were subsequently mounted onto Colorfrost Plus™ slides (Thermo Fisher 12-550-17) and dried overnight at 55ºC. Slides were maintained at room temperature in a desiccated environment until use.

### Human tissues and tissue processing

Human intestinal tissues were collected as previously described^15^ under the approval by the Health Science Research Ethics Board (HSREB) at the University of Western Ontario (HSREB-2597). Formalin fixed-paraffin-embedded tissue sections were prepared and stored at 4°C until immunofluorescence experiments.

### Immunofluorescence

Following deparaffinization and rehydration, Tris-EDTA Solution (EMS, Cat No. 64142-09) was used for antigen retrieval. Slides were submerged in 1X Tris-EDTA heated to 65°C and placed in the microwave for 14 minutes at power level 1. After cooling to room temperature, slides were permealized with 0.2% Triton X-100 for 30 minutes and blocked with 5% donkey serum in PBS for 4 hours. Mouse tissue sections were further blocked with donkey anti-mouse Fab fragments (Jackson Laboratories, Cat No. 715-007-003) at a 1:20 dilution for 1 hour. Primary antibodies for IL-25 (Invitrogen, Cat No. MA1-41067) and DCAMKL1 (Abcam, Cat No. ab31704) were diluted 1:100 and 1:200, respectively, with 2% donkey serum in PBS, and incubated with the pre-blocked slides overnight at 4°C in a humidified chamber.

The following day, slides were washed with 0.2% Triton X-100 in PBS and incubated with secondary antibodies donkey anti-mouse AF488 (Invitrogen, Cat No. A21202) and donkey anti-rabbit AF594 (Invitrogen, Cat No. A21207) at a dilution of 1:500 for 2 hours at room temperature. After subsequent washes, slides were stained with DAPI for 10 minutes. Coverslips were adhered and slides were imaged the following day. Brightfield and fluorescent images were acquired with the Olympus IX81 microscope and ImageJ was used to generate figures.

## Notes

### Competing Interest Statement

The authors have declared no competing interest.

### Funding

- Natural Sciences and Engineering Research Council: RGPIN-2022-04187

## References

1. Gerbe, F., Brulin, B., Makrini, L., Legraverend, C., & Jay, P. (2009). DCAMKL-1 Expression Identifies Tuft Cells Rather Than Stem Cells in the Adult Mouse Intestinal Epithelium. Gastroenterology, 137(6), 2179–2180. 10.1053/j.gastro.2009.06.072

2. Gerbe, F., van Es, J. H., Makrini, L., Brulin, B., Mellitzer, G., Robine, S., Romagnolo, B., Shroyer, N. F., Bourgaux, J.-F., Pignodel, C., Clevers, H., & Jay, P. (2011). Distinct ATOH1 and Neurog3 requirements define tuft cells as a new secretory cell type in the intestinal epithelium. The Journal of Cell Biology, 192(5), 767–780. 10.1083/jcb.201010127

3. von Moltke, J., Ji, M., Liang, H.-E., & Locksley, R. M. (2016). Tuft-cell-derived IL-25 regulates an intestinal ILC2-epithelial response circuit. Nature, 529(7585), 221–225. 10.1038/nature16161

4. Howitt, M. R., Lavoie, S., Michaud, M., Blum, A. M., Tran, S. V., Weinstock, J. V., Gallini, C. A., Redding, K., Margolskee, R. F., Osborne, L. C., Artis, D., & Garrett, W. S. (2016). Tuft cells, taste-chemosensory cells, orchestrate parasite type 2 immunity in the gut. Science (New York, N.Y.), 351(6279), 1329–1333. 10.1126/science.aaf1648

5. Gerbe, F., Sidot, E., Smyth, D. J., Ohmoto, M., Matsumoto, I., Dardalhon, V., Cesses, P., Garnier, L., Pouzolles, M., Brulin, B., Bruschi, M., Harcus, Y., Zimmermann, V. S., Taylor, N., Maizels, R. M., & Jay, P. (2016). Intestinal epithelial tuft cells initiate type 2 mucosal immunity to helminth parasites. Nature, 529(7585), 226–230. 10.1038/nature16527

6. Leppänen, J., Helminen, O., Huhta, H., Kauppila, J. H., Miinalainen, I., Ronkainen, V.-P., Saarnio, J., Lehenkari, P. P., & Karttunen, T. J. (2016). Doublecortin-like kinase 1-positive enterocyte – a new cell type in human intestine. APMIS, 124(11), 958–965. 10.1111/apm.12599

7. Huang, L., Bernink, J. H., Giladi, A., Krueger, D., van Son, G. J. F., Geurts, M. H., Busslinger, G., Lin, L., Begthel, H., Zandvliet, M., Buskens, C. J., Bemelman, W. A., López-Iglesias, C., Peters, P. J., & Clevers, H. (2024). Tuft cells act as regenerative stem cells in the human intestine. Nature, 634(8035), 929–935. 10.1038/s41586-024-07952-6

8. Billipp, T. E., Fung, C., Webeck, L. M., Sargent, D. B., Gologorsky, M. B., Chen, Z., McDaniel, M. M., Kasal, D. N., McGinty, J. W., Barrow, K. A., Rich, L. M., Barilli, A., Sabat, M., Debley, J. S., Wu, C., Myers, R., Howitt, M. R., & von Moltke, J. (2024). Tuft cell-derived acetylcholine promotes epithelial chloride secretion and intestinal helminth clearance. Immunity, 57(6), 1243-1259.e8. 10.1016/j.immuni.2024.03.023

9. Nadjsombati, M. S., McGinty, J. W., Lyons-Cohen, M. R., Jaffe, J. B., DiPeso, L., Schneider, C., Miller, C. N., Pollack, J. L., Gowda, G. N., Fontana, M. F., Erle, D. J., Anderson, M. S., Locksley, R. M., Raftery, D., & Moltke, J.von. (2018). Detection of succinate by intestinal tuft cells triggers a type 2 innate immune circuit. Immunity, 49(1), 33. 10.1016/j.immuni.2018.06.016

10. Mayassi, T., Li, C., Segerstolpe, Å., Brown, E. M., Weisberg, R., Nakata, T., Yano, H., Herbst, P., Artis, D., Graham, D. B., & Xavier, R. J. (2024). Spatially restricted immune and microbiota-driven adaptation of the gut. Nature, 636(8042), 447–456. 10.1038/s41586-024-08216-z

11. Zwick, R. K., Kasparek, P., Palikuqi, B., Viragova, S., Weichselbaum, L., McGinnis, C. S., McKinley, K. L., Rathnayake, A., Vaka, D., Nguyen, V., Trentesaux, C., Reyes, E., Gupta, A. R., Gartner, Z. J., Locksley, R. M., Gardner, J. M., Itzkovitz, S., Boffelli, D., & Klein, O. D. (2024). Epithelial zonation along the mouse and human small intestine defines five discrete metabolic domains. Nature Cell Biology, 26(2), 250–262. 10.1038/s41556-023-01337-z

12. Haber, A. L., Biton, M., Rogel, N., Herbst, R. H., Shekhar, K., Smillie, C., Burgin, G., Delorey, T. M., Howitt, M. R., Katz, Y., Tirosh, I., Beyaz, S., Dionne, D., Zhang, M., Raychowdhury, R., Garrett, W. S., Rozenblatt-Rosen, O., Shi, H. N., Yilmaz, O., … Regev, A. (2017). A single-cell survey of the small intestinal epithelium. Nature, 551(7680), 333–339. 10.1038/nature24489

13. Hickey, J. W., Becker, W. R., Nevins, S. A., Horning, A., Perez, A. E., Zhu, C., Zhu, B., Wei, B., Chiu, R., Chen, D. C., Cotter, D. L., Esplin, E. D., Weimer, A. K., Caraccio, C., Venkataraaman, V., Schürch, C. M., Black, S., Brbić, M., Cao, K., … Snyder, M. (2023). Organization of the human intestine at single-cell resolution. Nature, 619(7970), 572–584. 10.1038/s41586-023-05915-x

14. Bialkowska, A. B., Ghaleb, A. M., Nandan, M. O., & Yang, V. W. (2016). Improved Swiss-rolling Technique for Intestinal Tissue Preparation for Immunohistochemical and Immunofluorescent Analyses. Journal of Visualized Experiments: JoVE, (113), 54161. 10.3791/54161

15. Krishnamurthy M., Li J., Fellows G.F., Rosenberg L., Goodyer C.G., Wang R. (2011) Integrin {alpha}3, but not {beta}1, regulates islet cell survival and function via PI3 K/Akt signaling pathways. Endocrinology 152(2):424–435. doi:10.1210/en.2010-0877

